# Noise signature in added size suggests bacteria target a commitment size to enable division

**DOI:** 10.1101/2020.07.15.202879

**Authors:** César Nieto, Juan Arias-Castro, César Vargas-García, Carlos Sánchez, Juan Manuel Pedraza

**Affiliations:** Department of Physics, Universidad de los Andes, Bogotá, Colombia; Department of Systems Biology, Harvard Medical School, Boston, Massachusetts 02115, USA; AGROSAVIA Corporación Colombiana de Investigación Agropecuaria. Mosquera. Colombia

## Abstract

Recent experiments suggested that sizer-like division strategy, a deviation from the adder paradigm might be produced by additional degradation events of cell division machinery molecules. We revisited single cell size data from a recently microfluidics setup using the above model. We observed that such additional degradation process, although reproduces size observations in the mean sense, it is unable to capture cell size fluctuations. We further extended recently proposed power law models to include commitment size. Our proposal is in agreement of both mean and fluctuation profiles seen in experiments. Our approach suggests further uses of noise profiles on dissecting cell size regulatory mechanisms.

**SIGNIFICANCE:** We contrast cell division models against bacteria cell size data in minimal media. Our results seems to support the idea that division starts once bacteria meet a given commitment size.

---

To maintain tight distributions on size, bacteria must exert control over their division events (1, 2). Recent phenomenological data show that these bacteria often divide using the *Adder* strategy, where the size added during cell-cycle (birth to division) is uncorrelated with the size at birth (3, 4). Recent experiments and mathematical modeling have suggested size-dependent accumulation of FtsZ up to a critical threshold to be a putative biophysical mechanism resulting in the *Adder*. Significant deviations from this strategy occur in some slow-growing media, whereby the added size between birth to division is negatively correlated with the size at birth (also known as *sizer-like* strategy) (5, 6).

Three competing models explain these negative correlations: a) Degradation of FtsZ (5), b) nonlinear size-dependency of FtsZ’s accumulation rate, (7, 8) and c) existence of an additional size-control mechanism that allows division to proceed if a cell is at least a certain size (6, 9). Here we show that these models can be discriminated by the correlations between noise in the added size and size at birth. Specifically, mechanism a) has almost null correlation, b) has a small positive correlation, and c) has relatively high positive correlation. We present recent experimental data in agreement with c). We discuss possible molecular mechanisms behind this latter model and applications of this framework to more complex organisms.

## MODELLING DIVISION IN ROD-SHAPED BACTERIA

Consider cells growing exponentially in size *s* across time *t* with a growth rate *µ*, i.e.,

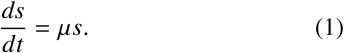

As depicted in Figure. 1, we assume that division triggers upon reaching a fixed number of steps *M* (10). In the classical *adder* strategy, the accumulation rate is proportional to the current cell size and the number of steps to division *M* can be estimated from the mean fluctuations in added size (5).

**Figure 1:**
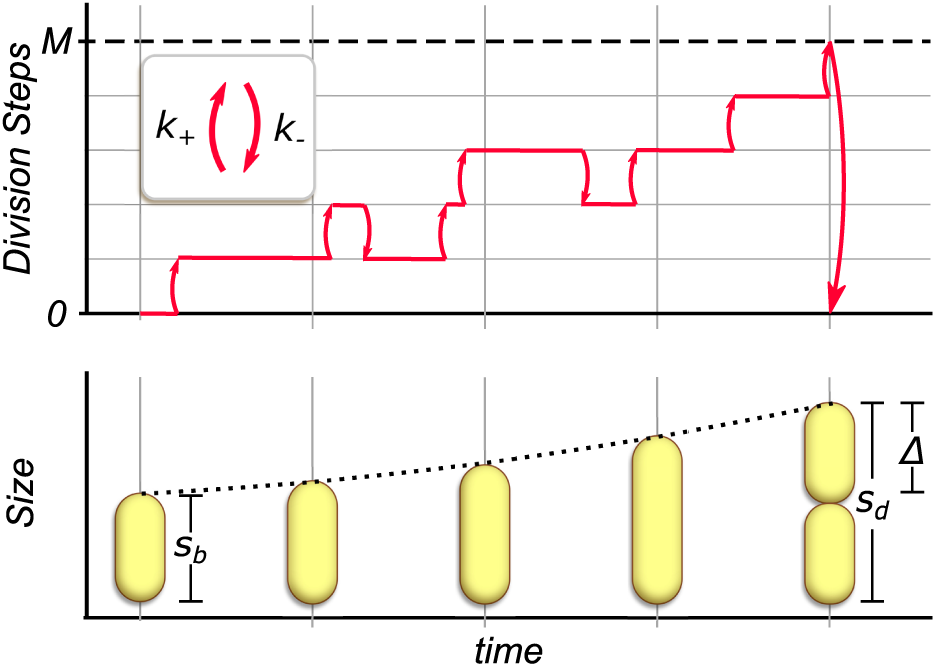
A general multistep model for the triggering of cell division. While bacteria grow exponentially (lower panel), division steps are performed at rate *k*_+_ and might revert at rate *k*_−_ (upper panel). Once a number of steps *M* is reached, the cell divides, the steps are reset to zero and the size is halved. The main variables of the bacterial cell cycle are also shown: the size at birth *s*_*b*_, the size at division *s*_*d*_ and the added size Δ = *s*_*d*_ − *s*_*b*_

Here, we will study how this model can be modified to obtain a division strategy with a negative correlation between added size and size at birth. In general, we will consider that each step happens at a rate *k*_+_ and, in some cases, might be reverted at rate *k*_−_. The size at step 0 and step *M* of the process correspond to the size at birth *s*_*b*_ and division *s*_*d*_, respectively as Δ = *s*_*d*_ − *s*_*b*_ is the size added between steps 0 and *M*.

Let *P*_*m*_(*t*) be the probability of *m* steps being completed time *t*. Given the rates *k*_+_ and *k*_−_, the dynamics of these probabilities are described by the master equation:

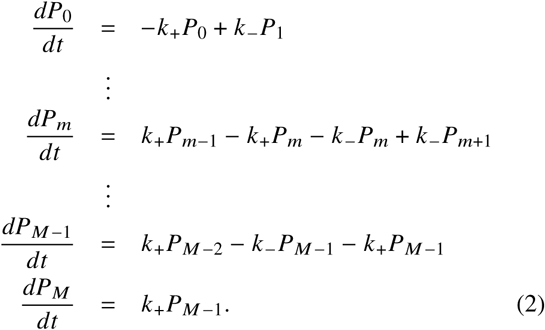

*P*_*M*_ is the probability of reaching the target steps *M* or equivalently, the probability of a division event to occur. Once the division happens, the process resets to step zero and size is halved. Using this *P*_*M*_(*t*) and the growth regime 1, the probability density *ρ*(*s*_*d*_) of size at division can be estimated as is shown in (5). The division strategy can be determined by observing the relationship between ⟨Δ ⟩ = ⟨*s*_*d*_⟩ − *s*_*b*_ and *s*_*b*_.

### The adder

The *Adder* strategy can be obtained if *k*_+_ and *k*_−_ are (5):

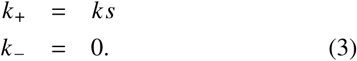

Assuming cells are growing exponentially and division process defined by both 2 and 3, the mean added cell size ⟨Δ⟩ is given by

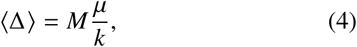

which is independent of the newborn cell size. This is the defining characteristic of the adder strategy (11).

Stochastic fluctuations (noise) in added size, quantified by the coefficient of variation 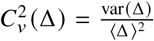, with var(Δ) being the variance of Δ, are given by:

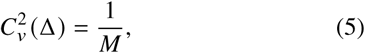

which is independent of the size at birth as observations confirm (5).

We now present the experiments performed to study the main properties of the *Sizer-like* in slow-growing bacteria, and later we introduce three possible mechanisms explaining this division strategy in terms of our proposed framework.

### Experiments

We revisited the experiments done in (5). These consist of time-lapse microscopy snapshots of long-term single-cell size dynamics of *Escherichia coli* bacteria (k12-MG1655). See (5) for method details. We studied 2740 cell-cycles of these bacteria growing in a *mother machine* micro-fluidic device (2) in M9 minimal media with Glycerol as carbon source.

Cell size was measured by the cell length given that width remains approximately constant trough the cell cycle. Comparison between experiments and theoretical results were done normalizing the cell size by 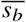 which corresponds to the average newborn size once bacteria have reached steady growth and division in both simulations and experiments.

### Sizer-like by a molecule that can be degraded

As (8) suggests, *Sizer-like* behavior can be obtained including active degradation of the division triggering molecule. In our framework, degradation is equivalent to a step backwards in the accumulation of the *M* events. In such case, the rates *k*_+_ and *k*_−_ are given by

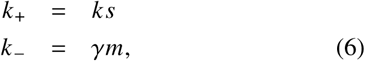

where *γ* is the rate per step at which division steps decrease.

As expected, a negative slope in Δ vs *s*_*b*_ as shown in Fig. S1 a., is obtained. The higher *γ*, the more pronounced is this slope. This model reduces to the adder when *γ* « *k* ⟨*s*⟩, with ⟨*s*⟩ being the mean cell size.

Once *γ* = 1.7*µ* is fitted from the trend line Δ vs *s*_*b*_, and *M* ≈ 18 is set from the mean noise 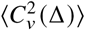, these fluctuations turn out to be relatively independent of size at birth in striking contrast with experiments (Fig S1.h.). To reproduce such dependency of the fluctuations, *γ* values would need to be large, increasing the slope in Δ vs *s*_*b*_ beyond what experiments suggest.

**Figure 2:**
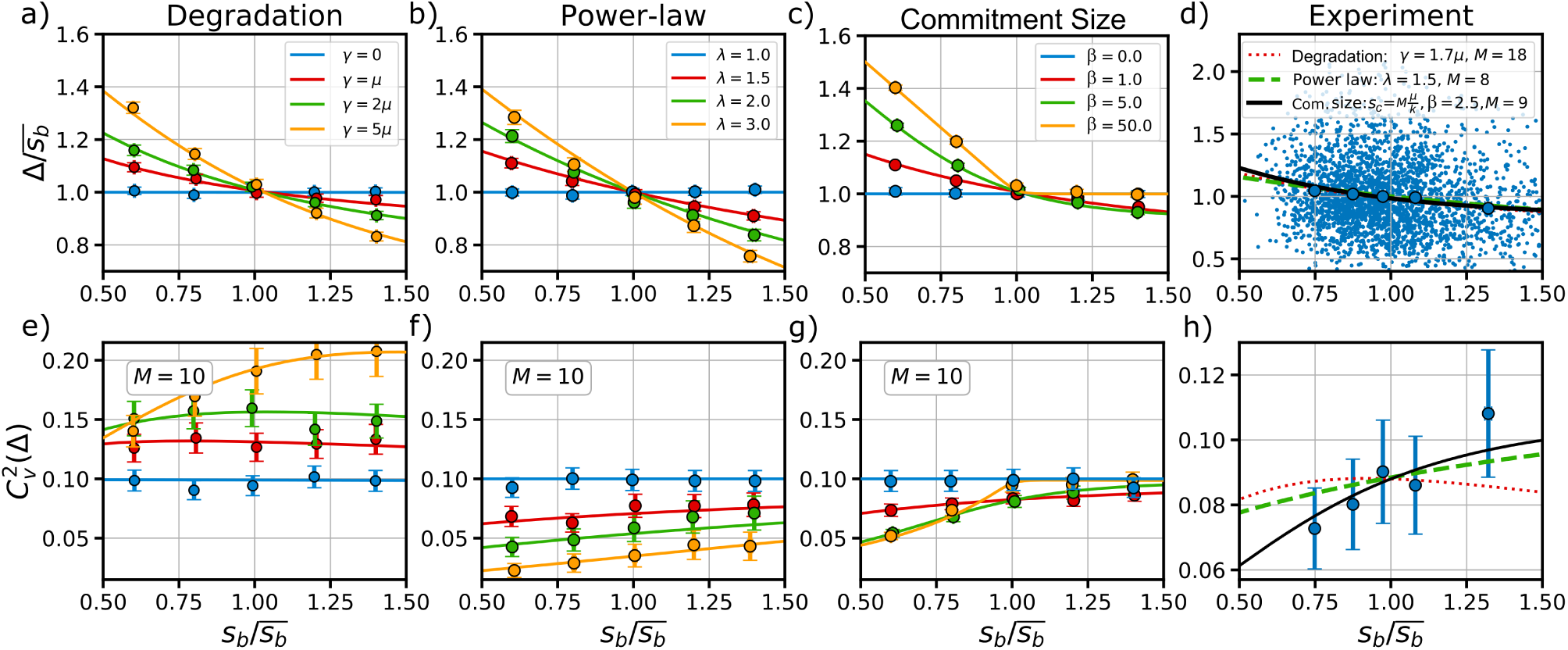
Trends on mean added size Δ and its stochastic fluctuations (noise) 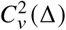 related to the size at birth *s*_*b*_ for respectively (a), (e) the model considering degradation for different values of *γ* in 6. (b), (f) the model considering a power law of the size for different exponents *λ* in 7. (c), (g) Division considering a initiation size with different Hill exponents *β* in 8 and 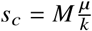. (d) (h) Measurements of division in *E. coli* growing in M9 minimal media with glycerol as Carbon source. Comparison between the three models and experiment is shown. Trend lines correspond to the numerical solutions of 2. Large dots are obtained from either simulations or experiments.

### Sizer-like by a molecule produced with rate proportional to a power of the size

An alternative mechanism, as per (5), suggests that a triggering molecule accumulates at rates proportional to a power *λ* of cell size, i.e,

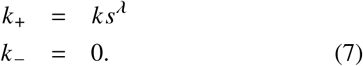

The process, defined by 2 and 7, reduces to the *Adder* strategy when *λ* = 1 resulting on a *Sizer-like* strategy when *λ >* 1 (Fig S1 b). The larger the *λ* the higher the slope in the division law.

The effect of this power law rate in the added size fluctuations 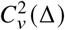 is two fold: it controls overall noise levels, and produces added size fluctuations that increase with the size at birth *s*_*b*_. After fitting the exponent (*λ* = 1.5) from the trend line on Δ vs *s*_*b*_ and *M* ≈ 8 from 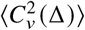, the resulting slope is rather modest compared with experiments (Fig S1 h).

### Sizer-like by introducing a commitment size

The experiments analyzed here suggest that cells with small size at birth have less fluctuations when compared with large newborns (5). Although the power law model reproduces qualitatively the above observation, it overestimate fluctuations for small newborns. Recent evidence suggests that cells may aim for a minimal size before starting division programs (6). Such minimal or commitment size *s*_*c*_ could be incorporated into our framework as per

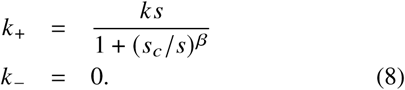

After fitting 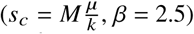 from the trend in Δ vs *s*_*b*_ and *M* = 9 from 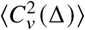, the expected dependence between 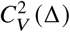 and *s*_*b*_, results in an increasing function with a slope higher than the predicted by the power law model.

To better understand the effects of this mechanism, two limits can be studied: For large *β* cell born with size lower that this commitment size *s*_*c*_, exhibit a perfect sizer strategy (Fig S1 c). Newborn cells with size higher than *s*_*c*_, exhibit an adder strategy (slope in Δ vs *s*_*b*_ is −1). This behavior is suggested by recent observations (12). On the other hand, when *β* is close to 0, the adder is obtained. For intermediate *β* values, as in our case, a relatively smoother trend line in Δ vs *s*_*b*_ is found.

## DISCUSSION

We compared recently proposed models that reproduce the *Sizer-like* behavior in slow growing *E. coli* bacteria in light of experimental data available from (5). As shown in Fig S1 d and h, all three models match the mean trend in added size vs size at birth. However, when noise in the added size 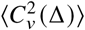 is plotted against size at birth *s*_*b*_, (see Supplementary information to get details of data analysis) these models show remarkable differences. While the model considering degradation predicts a nearly zero-slope relationship between added size noise and size at birth, the power-law model predicted an increasing function. The dependency shown by the latter seems too subtle to explain the fluctuation profile seen in experiments. The commitment model solves this discrepancy, predicting an increased dependency between size fluctuations and size at birth in agreement with experiments.

Recent experiments show different cell division strategies depending on whether they are in slow (*Sizer-like*) or fast (*adder*) growing media (6). In fast growing, cells are large and in slow growing, cells are small (2). The commitment size model reproduces such differences in division strategy as follows: in slow or poor media, average cell size falls below the commitment size threshold most of the time, disabling division process until the size requirement is met – by definition a sizer-like strategy. When cell is in fast or rich media, average size remains above the commitment size most of the cell cycle, enabling cell division at a rate proportional to the cell size, by definition an adder strategy.

How the commitment size model might be implement in bacteria is still in active debate. One alternative could be by tight synchronization between division and DNA replication processes, where division is enabled by meeting a target size required for DNA replication initiation (6). Alternatively, experimental evidence may support a division process mainly driven by a molecule or group of molecules that serve as a proxy of size within cells (13).

A mechanism to target size, in a similar way done by past studies (14), can be mediated by a molecule repressing the division initiation (See Supplementary Information). This molecule is produced in such a way that its number inside the cell remains constant, which means that its concentration is negatively correlated with the size (15). Given this repression, division starts once the molecule concentration is below some target. This idea reproduces both the trends in added size and in its noise.

This work shows how physiological processes at single cell level can be studied from their stochastic fluctuations – their noise signature – giving clues about the possible mechanisms behind these processes. This framework can be helpful in the design of future experiments where stochasticity is exploited fully as well as experiments studying complex cell cycles like those of yeast and other eukaryotic cells.

## AUTHOR CONTRIBUTIONS

C. N. Developed the model and analyzed the data, J.A.C. Performed the experiments, C.S. Made the image analysis, C.V.G. and J.M.P. guided the research. C.N, C.V.G. and J.M.P. wrote the article

## ACKNOWLEDGMENTS

We thank Johan Paulsson at Harvard Medical School for making his laboratory available to perform our experiments and image analysis. We also thank *Colciencias convocatoria para doctorados nacionales 647 de 2015* for the financial support. Finally we thank Khem Raj Ghusinga for the discussion.

## Supplementary Information

### GROUPING DATA TO OBTAIN LOCAL MOMENTS AS A FUNCTION OF ADDED SIZE

Experimental data was obtained by following individual cells in a Mother Machine, as specified in (5). As can be seen in Figure S1.a., it is not easy to visually compare the raw data with the trend line obtained from the numerical approach. To facilitate this comparison, the data was split in five quantiles for which the moments were obtained. Specifically, for the five points shown in Figure S1.a., the abscissa corresponds to the mean size at birth ⟨*s*_*b*_⟩ _*q*_ of each quantile and the ordinate is the average added size ⟨Δ⟩_*q*_ of such quantile. To display the noise signature, we plot the average variation of data 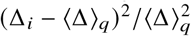 around each quantile average ⟨Δ⟩ _*q*_. The error bars correspond to a 95% confidence interval. This method was used for both experiment and simulation data presented in both Figure 2 in the main text and in Figure S1.

### A MOLECULAR MECHANISM TO TARGET A COMMITMENT SIZE

As a possible molecular implementation of the treshold mechanism mentioned in the main text, consider a case where division initiation is being repressed by an actively degraded protein *p* whose number is kept constant, with the repression described by a Hill function with reference value *p*_0_. The process can be described by the equations:

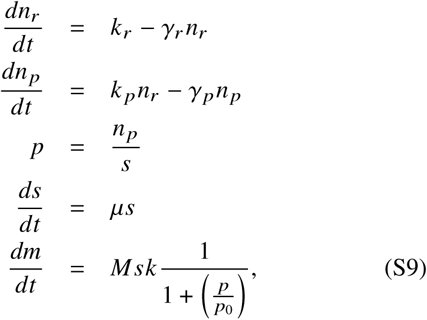

where *n*_*r*_ is the number of mRNAs for protein *p, n*_*p*_ is the number of such proteins and the cell size *s* grows exponentially at rate *µ*. We further assume, per (5), that multiple events *m* have to happen for division to be triggered. This could be an amount of protein, protofilaments or other chemical processes, which in the simplest case accumulate in proportion to the cell size *s* (5), but to implement the repression we propose an accumulation rate proportional to the cell size *s* times a Hill function dependent on the concentration of the repressor protein *p*. Once the number of events reaches the target number *M*, cells divide. The size is halved, the step number is reset to zero and proteins and mRNAs are segregated binomially. We let bacteria reach stationary distributions (approx 8 division times, each ln (2) /*µ* time units) and after reaching these stationary values, we used the next cell cycle to obtain the data in Figure S1.

The parameters used for this simulation are shown in table 1. The main difference with the phenomenological approach shown in Eq. 8 in the main text is that in the proposed molecular implementation the Hill function depends negatively on the concentration *p* of the repressor protein, which in turn depends negatively on the volume. In the phenomenological model all this is simplyfied to a single Hill-type activation..

**Table 1:** Parameters in the molecular mechanism.

| Parameter | Value |
| --- | --- |
| $\mu$ | $\ln(2)/\tau = \ln(2)$ |
| $k$ | $\mu$ |
| $\gamma_r$ | $5\mu$ |
| $k_r$ | $60\mu$ |
| $\gamma_p$ | $10\mu$ |
| $k_p$ | $110\mu$ |
| $p_0$ | 30 |
| $M$ | 10 |

---

**Figure S1:**
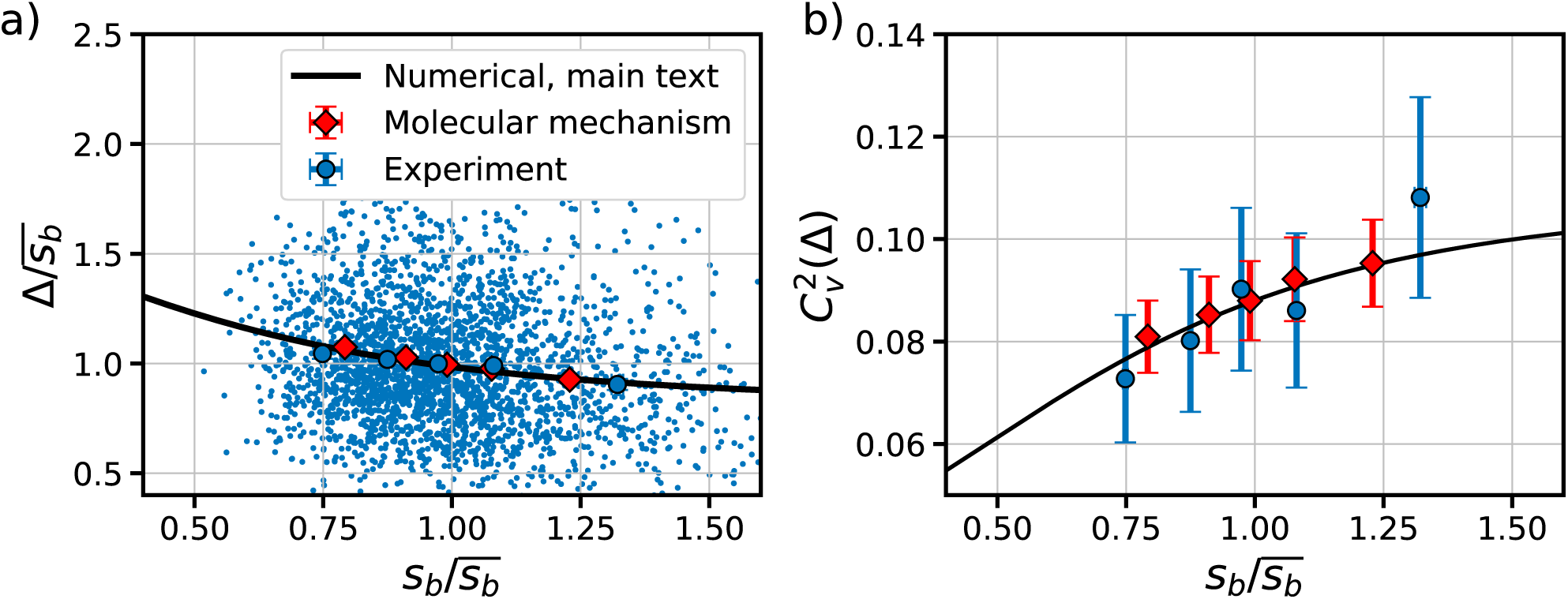
Comparison between the simulation of the molecular model (red diamonds), the phenomenological model (explained in the main text) (black line) and the experimental data (blue dots).

